# The role of the parietal lobe in task-irrelevant suppression during learning

**DOI:** 10.1101/2022.08.23.504946

**Authors:** F. Contò, S Tyler, P. Paletta, L. Battelli

## Abstract

**Background:** Attention optimizes the selection of visual information, while suppressing irrelevant visual input through cortical mechanisms that are still unclear. We set to investigate these processes using an attention task with an embedded to-be-ignored interfering visual input.

**Objective:** We delivered electrical stimulation to attention-related brain areas to modulate these facilitatory/inhibitory attentional mechanisms. We asked whether training on a task while being simultaneously exposed to visual features from a different task tested at baseline and post-training might influence performance on the baseline task.

**Methods:** In Experiment one, subjects performed an orientation discrimination (OD) task using a pair of gratings presented at individual’s psychophysical threshold. We then trained participants over multiple sessions on a temporal order judgment task (TOJ), using the exact same gratings but presented with different time offsets. On the last session we re-tested OD. We coupled training with transcranial random noise stimulation (tRNS) over the parietal cortex, the human middle temporal area or sham, in three separate groups. In Experiment two, subjects performed the same OD task at baseline and post-training, while tRNS was delivered at rest during training.

**Results:** Results showed that tRNS over parietal cortex facilitated learning of the trained task. Moreover, we found a *detrimental* effect on OD when subjects received parietal tRNS coupled with training, but a *benefit* on OD when subjects received stimulation while at rest.

**Conclusions:** These results clearly indicate that task-irrelevant information is actively suppressed during learning, and that this prioritization mechanism of selection likely resides in the parietal cortex.

## Introduction

Our visual environment is cluttered, yet we still perceive the visual scene effortlessly. To overcome our limited capacity to process multiple stimuli, there must be mechanisms that selectively direct our attention towards crucial visual targets, while inhibiting irrelevant information [1]. One way to study visual selection mechanisms is perceptual learning, the long-term improvement in performance on a visual task that results from practice [2]. Attention facilitates learning ([3]–[5], however whether it acts as a gating mechanism or through facilitation/inhibition of information is still unclear [6]–[8]. Several studies indicate that attention modifies spontaneous cortical activity to support behavior (e.g., [9]–[11]). In particular, endogenous attention, driven by a person’s expectations, exerts a top-down control, leading to a fast improvement in performance. Interestingly, many studies showed that the magnitude of perceptual learning increases with practice through filtering task-irrelevant information ([12]–[14]), likely mediated by voluntary top-down attention leading to enhanced selection of relevant information ([15], [16]). Top-down attention might thus induce a biased neural state: it enhances neural activity related to relevant information, whilst suppressing the neural response to irrelevant information ([17]). While attention can flexibly drive the enhancement of relevant visual stimuli, unattended visual information can still influence learning, as shown in task-irrelevant perceptual learning studies ([19], [20]). Although much is known about how voluntary attention can facilitate processing of task-relevant information (e.g., [21], [22]), less is known about facilitatory/inhibitory attentional mechanisms. One way to selectively enhance or inhibit cortical mechanisms of learning is transcranial electrical stimulation (tES) ([23]–[27]). tES delivers a weak electrical current to the brain through conductive electrodes placed on the scalp, and it changes the neuronal state depending on the polarity, intensity and frequency of stimulation ([28]). The induced neuromodulatory effects can outlast the stimulation ([29], [30]) and spread across task-related cortical networks ([31], [25], [32]). High-frequency transcranial random noise stimulation (tRNS), in particular, delivers a randomly alternating current to the brain, exerting a stochastic resonance effect, whereby it enhances the signal-to-noise ratio between cells firing in response to crucial targets relative to non-relevant noise information ([33]–[37]). Notably, several experiments have shown how training coupled with tRNS over task-related cortical areas can greatly benefit learning ([23]–[25], [38]).

Here we used tRNS to study the role of the parietal cortex in learning a task, while subjects are simultaneously exposed to a to-be-ignored task-irrelevant visual feature embedded in the stimuli. We questioned whether short sessions of tRNS to attention-related brain areas could induce rapid learning by modulating the facilitatory/inhibitory attentional mechanisms. We found that processing a target feature of a stimulus actively suppresses the processing of the task-irrelevant feature embedded in the stimulus, and that tRNS might act by inhibiting interfering information or enhancing target features by bringing them above threshold. We also studied the effect of stimulation alone with no training, to understand its potential to optimize cortical response in absence of behavioral training.

## Methods & materials

### Participants

Eighty-four neurologically healthy subjects (59 females, age range 18-35 years old), with normal to corrected-to-normal vision participated in the study. All subjects met all brain stimulation screening criteria and provided written informed consent approved by the ethics committee of the University of Trento. Subjects received monetary compensation for their participation in the experiment. 43 participants were recruited for Experiment 1, while 41 participants were recruited for Experiment 2; a total of 3 participants were detected as outliers and removed from statistical analysis (leaving 42 for experiment 1 and 39 for experiment 2).

### Experiment 1

#### Experimental Procedure

Subjects participated in a multi-session experiment (Figure 1A) that consisted of one pre-test session, three training sessions, and one post-test session (one session per day, for a total of 5 consecutive sessions). On Day 1, using a 3-1 staircase procedure subjects were tested on an Orientation Discrimination (OD) task where we measured their point of subjective equality (orientation values were adjusted to yield 50% accuracy in the task). Subjects then completed an OD task at threshold (pre-test) during which they were asked to determine whether two pair of gratings (presented to the left and right of fixation) had same or different orientation, thus judging the spatial orientation of the two stimuli (Figure1B). Subjects also performed the OD task at the end of the experiment, on Day 5 (post-test). On Days 2-4, subjects underwent three training sessions (one/day) on a temporal order judgment task (TOJ: Figure1C) during which participants were asked to indicate whether the left or the right grating appeared first, thus judging the temporal order of the two stimuli. Using the method of constant stimuli, the two target stimuli were presented with varying temporal offset asynchronies. Importantly, on each trial, the orientation information at individual threshold was embedded in the visual stimuli, and tRNS was delivered concurrent with training. Subjects were randomly assigned to one of three stimulation conditions: bilateral tRNS over parietal cortex (Parietal group), bilateral tRNS over middle temporal cortex (hMT+ group), and sham stimulation (Sham group).

**Figure 1.**
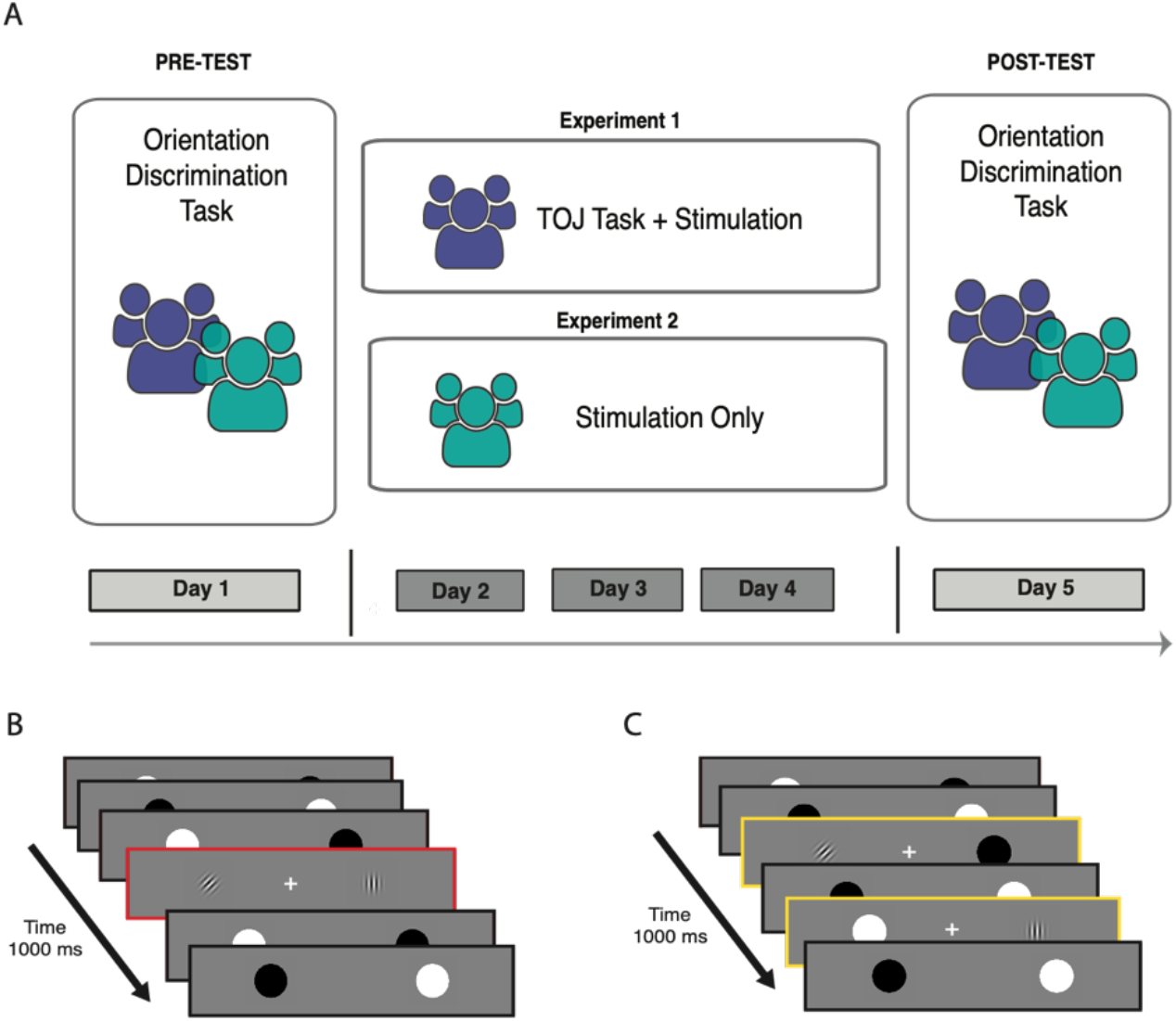
Experimental timeline for Experiment 1 and Experiment 2, and Tasks. **A)** Subjects of both experiments (Experiment 1 and Experiment 2) performed the Orientation Discrimination task on the pre-test and post-test session (Day 1 and Day 5). On days 2 to 4, subjects included in Experiment 1 underwent training on the Temporal Order Judgment Task while receiving stimulation, while subjects included in Experiment 2 received stimulation only. B) In the Orientation Discrimination Task, subjects had to discriminate the orientation of the two Gabors, embedded in a flicker cycle. C) In the Temporal order Judgment task, subjects had to determine which of the two Gabors appeared first. The two Gabors were presented with orientation values tilted at the individually measured threshold.

### Trial sequence and Stimuli characteristics

For both tasks, OD and TOJ, each trial consisted of a pair of Gabor discs (each 1.6 degree of visual angle), positioned 4° to the left and right of a central fixation cross. Subjects were instructed to maintain fixation on the central cross for the entire duration of the experiment. At the beginning and at the end of each trial the two discs appeared as stationary texture pattern for 500ms, which was used to mask any cue and prevent any resulting perceptual afterimages from the final frame. In-between the stationary states, the two discs flickered at a fixed 7.5Hz between high-contrast black and white for 1000ms. The flicker rate was set based on previous research that indicated to be well within subjects’ temporal discrimination threshold ([39], [40]). During the flicker presentation, the two discs appeared as Gabor patches for 133ms, after which the texture mask was presented again. At the end of the second texture mask, the two discs disappeared from the screen and participants were asked to report whether the two Gabors had same or different orientation for the OD task, or which of the two Gabors appeared first in the TOJ task. A visual feedback on the screen indicated whether the response was ‘Incorrect’ or ‘Correct’.

All Gabors were presented with a spatial frequency of 3 cycles/deg and reduced 50% contrast against a uniform gray background (47.5 cd/m!), with a fixation cross positioned in the center of the screen for the entire duration of the trial (Figure 1B and 1C). In the OD task, one of the two Gabors was tilted along the vertical line with a degree that corresponded to the 50% individual threshold calculated with the staircase procedure at the beginning of the experiment. In the TOJ task, the two Gabors were presented with a range of fixed temporal offset asynchronies (+/− 75, 41.66, 25, 58.33ms). In addition, one of the two discs was tilted along the vertical line with a degree that corresponded to the 50% individual orientation threshold calculated for the OD. The offset feature (task-relevant feature) and the orientation feature (task-irrelevant feature) were both presented counterbalanced per visual field. Stimuli were displayed on a 22-in. LCD monitor with a 120 Hz refresh rate controlled by a DELL computer equipped with Matlab (The MathWorks, Natick, MA) and Psychtoolbox 3.0.8 ([41], [42]). Participants were seated 57 cm from the screen in a dark and quiet room and used a chin rest to ensure consistent positioning.

### Stimulation Procedure

All participants were provided with a short introduction to brain stimulation and safety information. A battery-driven stimulator (DC-Stimulator, NeuroConn, Ilmenau, Germany) was used for electrical stimulation. For the two active high-frequency tRNS conditions (Parietal and hMT+), 2mA current was applied for 25 consecutive minutes with random alternating frequency delivered between 101 and 640 Hz. For sham stimulation, the machine was turned off after the fade-in phase. Two rubber electrodes (size=5×7 cm), contained in sponges soaked in saline solution, were placed on the subject’s head and were kept fixed on the stimulation sites with a rubber band and head cap. Stimulation sites were identified using the International electroencephalographic 10/20 system for scalp electrode localization. The center of the saline-soaked electrode was placed bilaterally over PO7/PO8 for hMT+ condition and over P3/P4 for parietal and sham conditions. Modeling of the electric field following stimulation was performed with free SimNIBS ([43], [44]) and visualized separately for the two active stimulation condition (Figure 2B and C).

**Figure 2.**
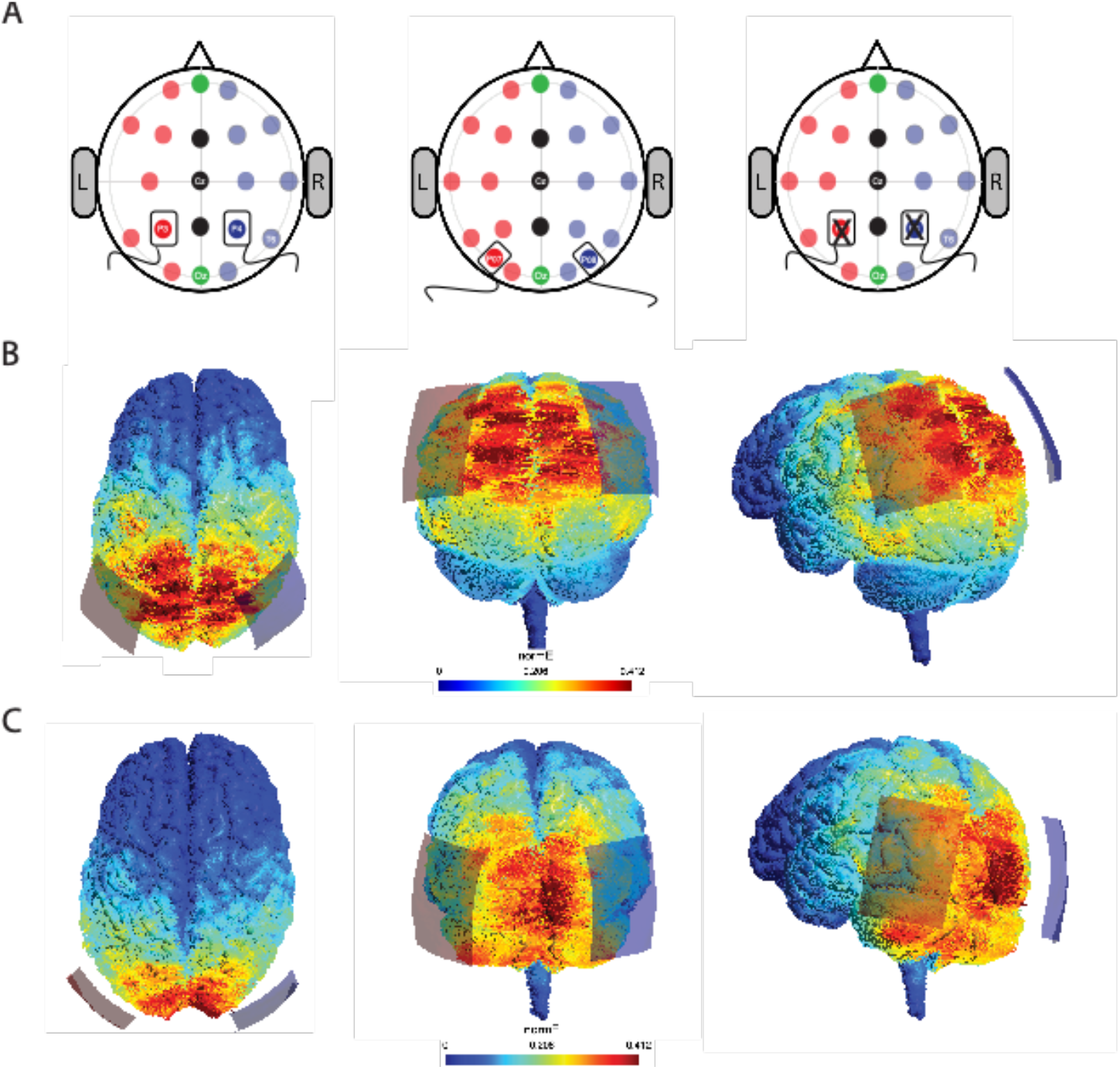
Stimulation Settings and Electric field distribution after stimulation. A) Stimulation sites were localized using EEG 10/20 system. Saline-soaked electrodes were placed over P3 and P4 for bilateral parietal and sham stimulation, and over P07 and P08 for hMT+ stimulation. Electric field distribution on the brain induced by Parietal and by hMT stimulation (Panel B and C, respectively).

### Experiment 2

The experimental procedure of Experiment 2 was the same as Experiment 1, except that subjects included in this second experiment did not undergo the three training sessions on the second task (TOJ), but received offline tRNS stimulation only (Figure 1A). During Days 2-4, subjects included in Experiment 2 received stimulation with the same randomly assigned stimulation conditions (Parietal, hMT+ and Sham) while resting (subjects were asked to sit quietly in a semi-dark room and not do anything while receiving stimulation).

## Results

### Experiment 1

We first measured the effect of stimulation combined with training on the OD task, which subjects performed on the pre- and post-test sessions. Next, we evaluated the effects of stimulation during the training sessions on the TOJ task.

#### Orientation Discrimination

Performance in the orientation discrimination task was quantified as the mean percentage of correct responses. Pre- and post-test accuracy is shown in Figure 3, separately for the three stimulation groups. First, to control that the three stimulation groups did not differ at baseline, prior to any manipulation, we compared performance on the pre-test session (Day 1) by performing a one-way ANOVA. This analysis revealed that the three groups did not differ significantly (F(2,39)=.635, p=.535, ηp2=.032). We next analyzed whether the combination of training and stimulation impacted subjects’ performance on the OD task by performing a mixed repeated measures ANOVA, with session as within-subjects (pre versus post) and stimulation group as the between-subjects factor (Parietal, hMT+, sham). These analyses revealed a strong significant interaction between stimulation condition and day of performance (F(2,39)=14.433, p<.001, ηp2=.425), indicating that accuracy scores of the pre-and post-test sessions differed significantly between stimulation groups. We found a significant main effect of stimulation condition alone (F(2,39)=4.743, p=.014, ηp2=.196), and no main effect of session on OD (F(1,39)=3.939, p =0.054, ηp2=.092). Given the significant interaction found in the main mixed repeated measure ANOVA, an analysis of simple main effects for stimulation factor was also performed for the post-test session. This analysis revealed a simple main effect of stimulation (F(2,39)=8.85, p<=.001, ηp2=.312). Post-hoc pairwise comparisons between post-test performances, showed that mean accuracy was significantly different between hMT+ and parietal condition (p<.001, Cohens’ d=1.565, Bonferroni p-corrected=<.001,) and between parietal vs sham (p=.045, Cohens’ d=−0.815, Bonferroni p-corrected=.135), but not between hMT+ and sham (p=.0501, Cohens’ d=.750, Bonferroni p-corrected=.150). In a second set of pairwise comparison between the pre- and the post-test sessions (Day1 and Day5), separated by stimulation factor, we found that hMT+ and sham group did not differ significantly from pre- to post-sessions (p=.225, Cohens’ d =−.604; p=.767, Cohens’ d=.124 respectively), while a significant difference was found for the parietal group (p=.007, Cohens’ d=1.620). These comparisons indicated that only performance of the parietal group was significantly different between the pre- and post-session (accuracy pre-test *M*=54.8% correct, σ=.9.643; accuracy post-test *M*=42.5% correct, σ=12.592).

**Figure 3.**
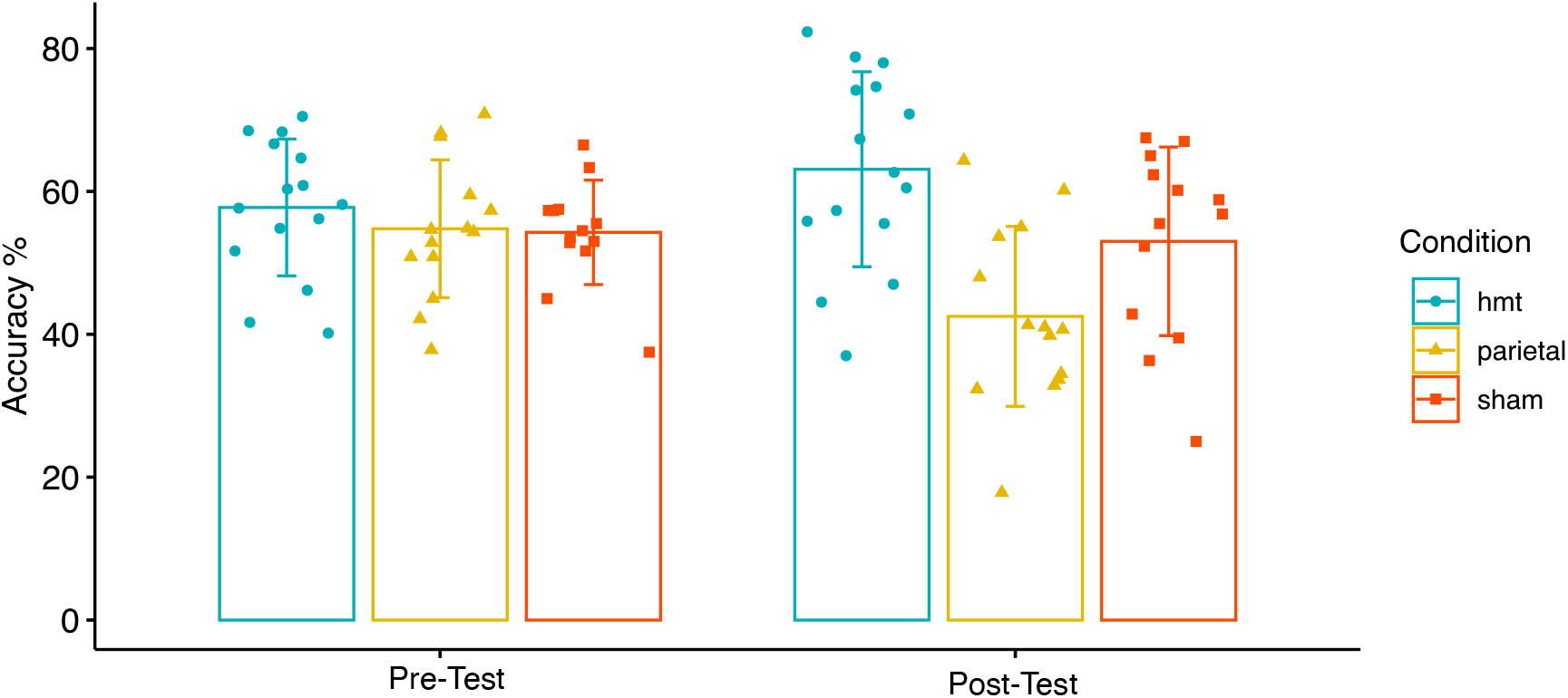
Pre-test and post-test performance. OD performances measured before and after stimulation and training procedure are shown for the three conditions, separately (blue for hMT+, yellow for Parietal and red for Sham). Vertical bars indicate standard deviation. Blue dots, yellow triangles and red squares represent individual subject’s data.

#### Temporal Order Judgment

We analyzed the time course of learning on the TOJ task across the three training sessions. To measure changes in performance over time, we calculated the percent improvement across the training session by calculating the difference between the first and the last training session (∆ Total Training Improvement=Day4-Day2), thus obtaining the mean normalized performance (delta values) for each subject and each group. Changes in performance between Day4 and Day2 are depicted in Figure 4. We then performed a one-way ANOVA to test whether and when there was a significant difference between performance improvements across conditions. We found a significant difference between the three stimulation groups (F(2,36)=4.617, p=.016, ηp2=.204; Figure 4). Post-hoc test comparisons indicated that there was a significant difference between the total performance improvement measured for the parietal relative to hMT+ group (p=.018, Bonferroni corrected), while the difference between parietal and sham (p=.106, Bonferroni corrected), and between hMT+ and sham was not significant (p=1, Bonferroni corrected).

**Figure 4.**
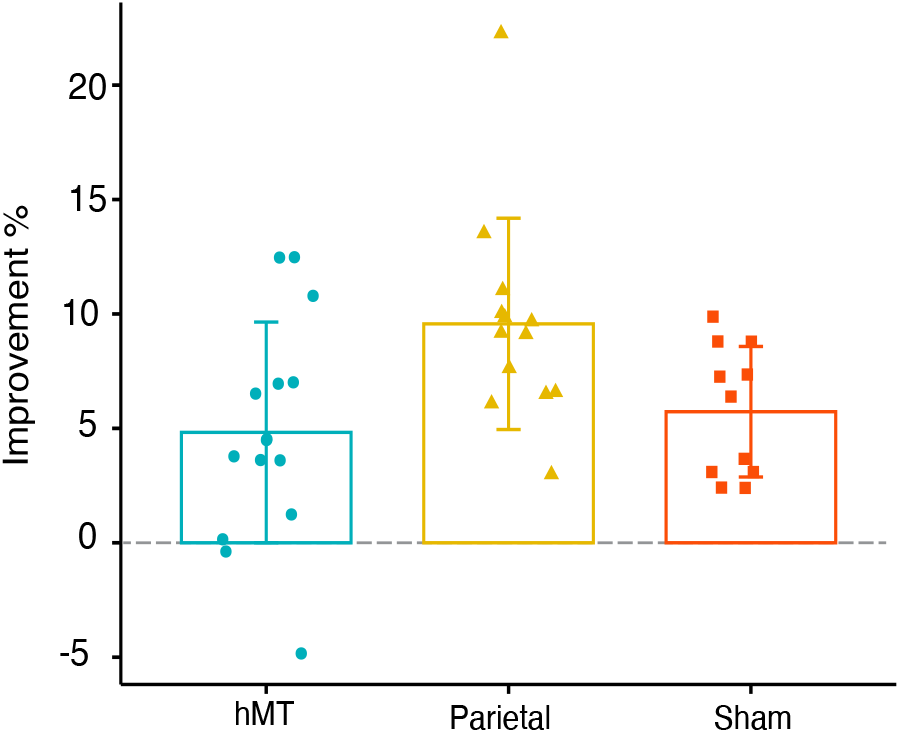
Performance change in the TOJ task. Delta values corresponding to the total difference in performance between training sessions (last training day relative to the first training day) are displayed per stimulation condition (hMT+, Parietal, Sham). Individual performance change is represented as blue dots, yellow triangles and red squares for hMT+, Parietal and Sham condition, respectively. The horizontal dotted line represents zero change in performance. Error bars show standard deviation.

### Experiment 2

We measured the effects of stimulation on the OD task, which was performed on the pre- and post-test sessions, with the 3-day stimulation-at-rest sessions. As for Experiment 1, performance was quantified as the mean percentage of correct responses. Accuracy measured in the pre- and post-test is shown in Figure 5, separately for the three stimulation groups. The effect of multi-session offline stimulation on OD performance was analyzed by performing a mixed repeated measures ANOVA, with session as the within-subjects factor (Pre versus Post) and stimulation as between-subjects factor (Parietal, hMT+, sham). We found a strong significant interaction between stimulation condition and day of performance (F(2,36)=11.510, p<.001, ηp2=.390), which indicates that accuracy scores of the pre- and post-test sessions significantly differed depending on the stimulation group. No main effect of stimulation condition alone was found (F(2,36)=1.269, p=0.29, ηp2=.066), while we found a significant effect of time on OD (F(1,36)=12.474, p=.001, ηp2=.257). We then performed a one-way ANOVA to compare the performance of the three stimulation groups on the pre-test session, prior to stimulation, and we found no statistically significant effect (F(2,36)=.881, p=.881, ηp2=.007), indicating that the three stimulation groups did not differ at baseline. To further analyze the significant interaction found in the mixed repeated-measures ANOVA, we performed an analysis of simple main effects for stimulation factor on post-test-session performance. A one-way ANOVA revealed a significant effect of stimulation (F(2,36)=3.36, p=.046, ηp2=.157), indicating that performance differed between stimulation groups on the post-test session. Next, post-hoc pairwise comparisons between post-test performances found that the mean accuracy score was significantly different between parietal and sham groups (p=.0138, Bonferroni p-corrected=.0413; Cohen’s d=.943,), but not between hMT+ and parietal (p=.248, Bonferroni p-corrected=.743; Cohen’s d=−0.489) nor between hMT+ and sham (p=.189, Bonferroni p-corrected=.568; Cohen’s d=.541)

**Figure 5.**
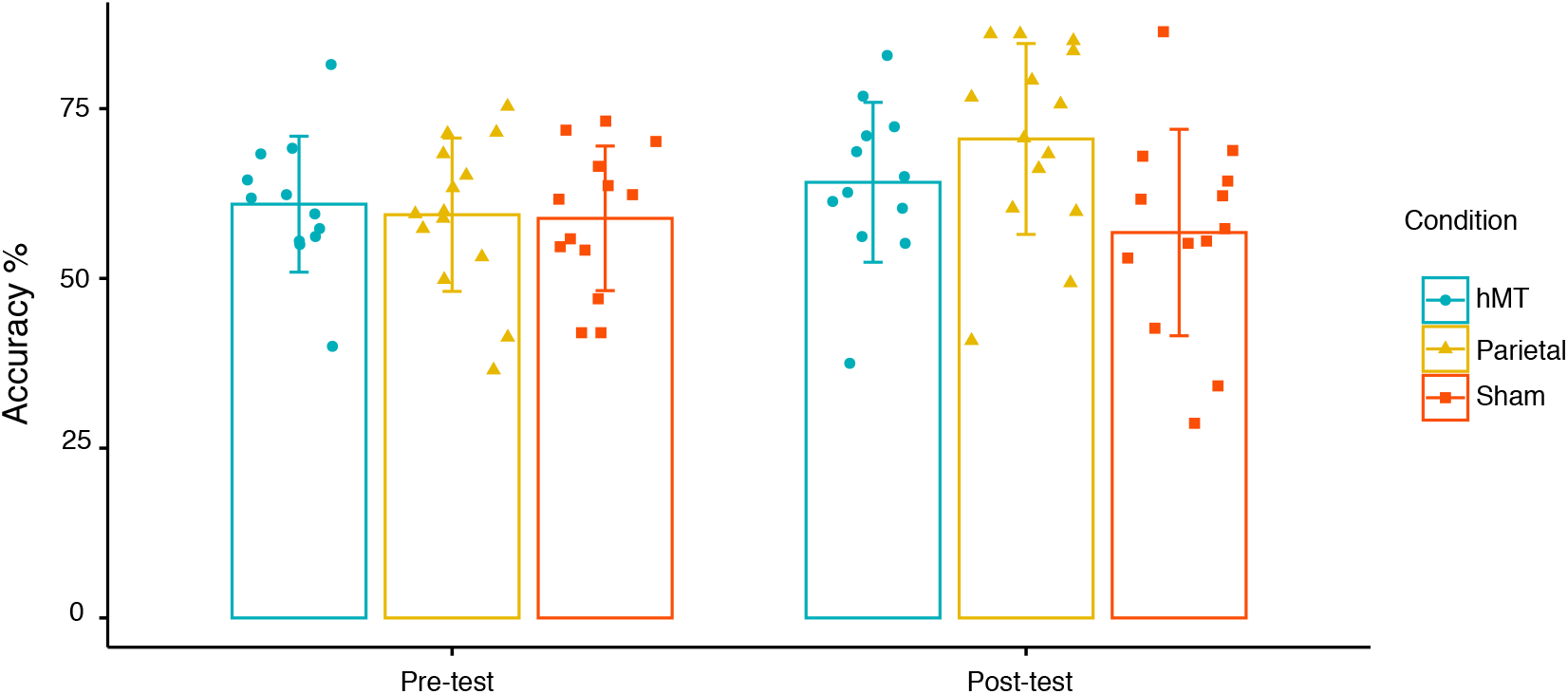
Pre-test and post-test performance. OD performances measured before and after stimulation and training procedure are shown for the three conditions, separately. Vertical bars indicate standard deviation. Blue dots, yellow triangles and red squares represent individual subject’s data.

Next, we run a second set of post-hoc pairwise comparison between the pre- and the post-test sessions, organized by stimulation factor (we compared accuracy scores between pre- and post-test session, for each stimulation condition). We found that performance of the parietal group differed between pre- and post-test (p=.0288, Cohens’ d=−2.120), while the performance of hMT+ and sham groups did not differ significantly (p=.478, Cohen’s d=−.477; p=.688, Cohens’d=.227 respectively).

#### Comparing OD in Experiment 1 and 2

In a new set of analyses, we compared the performance change in OD found in Experiment 1 relative to Experiment 2. Hence, we analyzed the effects of tRNS combined with task-irrelevant training (training coupled stimulation) vs no training (off-line stimulation) on the OD task performed by all subjects on the pre- and post-test sessions. We evaluated whether the combination of training vs no training and the stimulation factors impacted subjects’ performance on the OD feature. The learning effect was measured by calculating the percent improvement difference between the pre-test and the post-test session, thus obtaining the normalized mean value, which represents the total difference in performance between the two sessions per group (∆Performance|Improvement = Post-Test Performance – Pre-TestPerformance). We performed a two-way ANOVA to test whether there was a significant difference between performance improvements across groups, examining the effects of stimulation (Parietal, hMT+, sham) and training group (tRNS+task, tRNS only) on the delta scores. The analysis revealed a statistically significant interaction between training group and stimulation condition (F(2,75) = 21.482, p <=.001, ηp2=.364). Finally, training (F(1,75)=14.180, p<=.001, ηp2=.159) and stimulation factor (F(2,75)=4.841, p=0.022, ηp2=.097) were also statistically significant.

Given the statistically significant interaction found between stimulation condition and procedure (training vs no training coupled with stimulation), we performed an analysis of simple main effects for stimulation. There was a statistically significant difference in mean scores for subjects who underwent the training coupled with stimulation procedure (F(2, 75) = 17.2, p=<.001, ηp2=.390) as well as for subjects who underwent the offline stimulation procedure (F(2, 75) = 9.10, p=<.001, ηp2=.425) depending on the stimulation condition (hMT+, parietal or sham). These analyses revealed that the stimulation significantly affected delta scores of subjects included in Experiment 1 as well as in Experiment 2.

Next, to further investigate the significant interaction effect, we ran a series of pairwise t-test comparisons. The first set of pairwise comparisons analyzed the different performance between stimulation groups organized by procedure factor (we compared each combination of stimulation condition, for each procedure factor; significant results are depicted in Figure 6). Behavioral performance was significantly different between parietal and sham groups (p<.001, Bonferroni p-adjusted<.001; Cohen’s d= 1.784), and between parietal and hMT+ (p=.008, Bonferroni p-adjusted=.0251; Cohen’s d=−1.323) for the offline stimulation condition of Experiment 2, as well as between parietal and hMT+ (p=<.001, Bonferroni p-adjusted=<.001; Cohen’s d=2.133), and between parietal and sham (p=.002, Bonferroni p-adjusted=.007; Cohen’s d=−1.236) in Experiment 1. The second set of pairwise comparisons were run between Experiment 1 and 2, organized by stimulation condition. We found a significant difference for the two parietal stimulation conditions, with the parietal group of Experiment 1 showing a *decrease* in performance, while the parietal group of Experiment 2 showed a *boost* in performance (p<.001), while no significant differences were found between two procedures groups that underwent either hMT+ (p=.499) or sham stimulation (p=.828).

**Figure 6.**
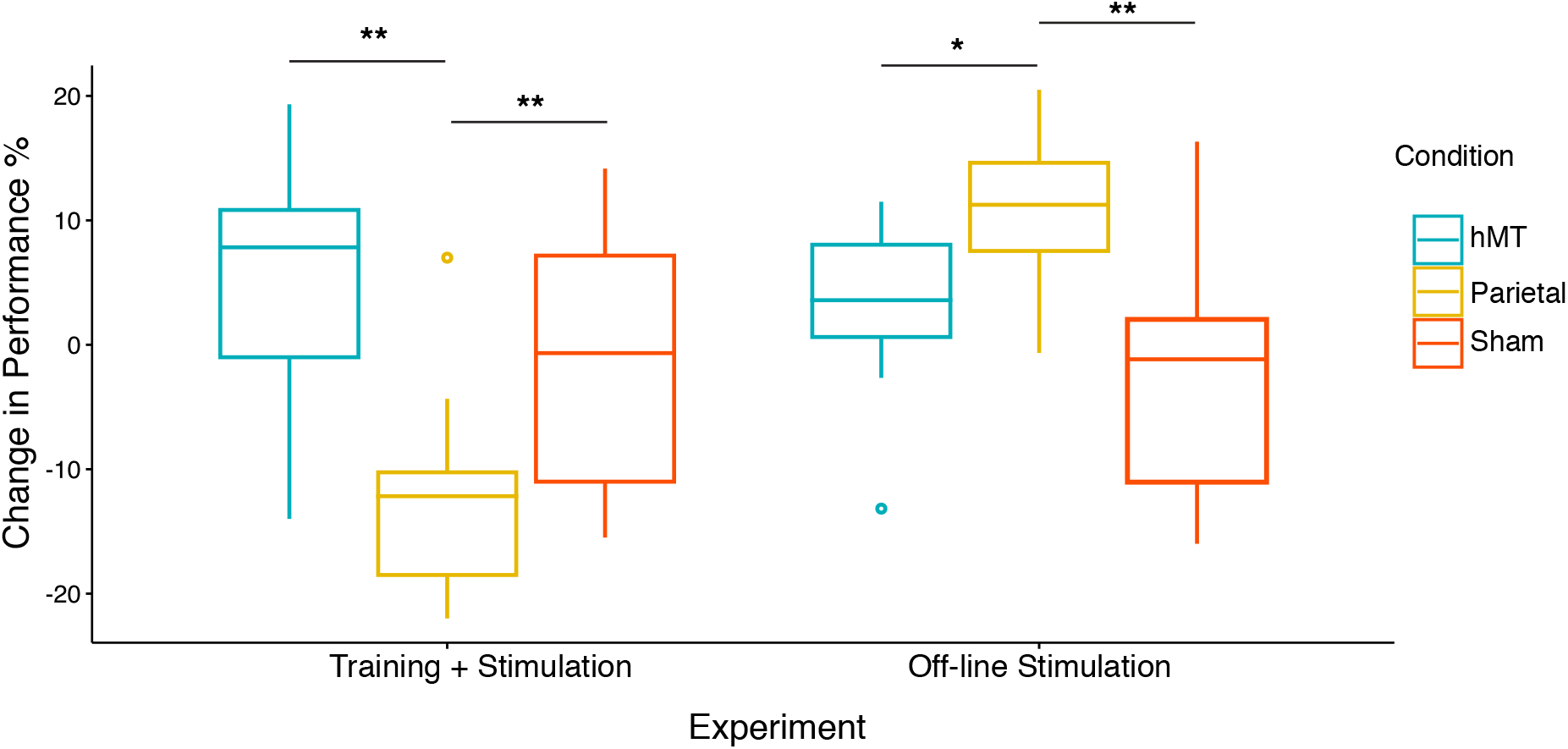
Behavioral difference between the pre- and the post-test sessions. Box plots for normalized mean performance represented per stimulation condition (green for hMT+, orange for Parietal and purple for Sham) and procedure (training coupled with stimulation vs off-line stimulation) on the OD task. The center line in the middle of the box is the median of each data distribution, while the box represents the interquartile range (IQR) with the lower quartile representing the 25^th^ percentile and the upper quartile representing the 75^th^ percentile. Asterisks represent significant comparisons (*=<0.05, **=<0.001).

## Discussion

We used noninvasive neuromodulation to study the causal role of the parietal cortex in facilitatory and inhibitory mechanisms of attentional processes during perceptual learning. In the first experiment we tested whether a task-irrelevant stimulus feature interfered with learning of a relevant stimulus, and we used tRNS to directly manipulate the rate of learning. We then analyzed subjects’ performance change in a task-irrelevant visuospatial task (OD task) whose visual feature was embedded in the task-relevant stimuli used for the training (TOJ task). Performance in the OD task was tested at baseline and at the end of the training. We found that when subjects trained on a task while concurrently receiving tRNS bilaterally over the parietal cortex (Experiment 1), their performance in the task-irrelevant (embedded) feature was strongly *impaired*. However, when subjects received stimulation over the parietal cortex without any perceptual training (Experiment 2), their performance in the OD task *improved*. This indicates that neuromodulation coupled with training resulted in an efficient suppression of concurrently presented, yet task-irrelevant, visual features. The interaction among multiple stimulus features (e.g., orientation vs temporal features) can be interpreted as a competition for neural processing. Perceptual learning has been hypothesized to work by selecting and/or strengthening crucial neural channels while reducing input from irrelevant channels, resulting in greater task-relevant visual information extraction ([12], [45]). Moreover, animal ([46]) and human studies ([47], [48]) demonstrated that attentional mechanisms inhibit distracting information by enhancing behaviorally relevant stimuli during practice ([49], [50]). Our neuromodulatory procedure might have affected these mechanisms, by facilitating the inhibitory processes to prioritize the relevant features during learning.

Our findings further suggest that learning is mediated by selective and inhibitory attentional mechanisms, and that parietal stimulation can impact these processes. The location of our stimulation, P3 and P4 of the 10-20 EEG systems ([51], [52]), roughly corresponding to the intraparietal sulcus (IPS), have been demonstrated to be directly involved in visuospatial attention ([53]), and our results indicate their crucial role played in actively suppressing irrelevant information while prioritizing relevant ones. Our stimulation procedure might have strengthened these dual attentional processes that underlie the emergence of learning of relevant visual information ([12], [13]). We also inspected the effect of the stimulation on the TOJ task used during training (Day2 to Day4). We analyzed the overall behavioral gain by contrasting performance of the last day normalized to the first day of training, and we found that subjects that received parietal tRNS significantly improved compared to hMT+ and sham groups. These results support the crucial role of the parietal lobe in driving a rapid improvement in attention-mediated perceptual learning within few training sessions. The importance of the parietal cortex in temporal attention task has been extensively demonstrated in previous neuropsychological and stimulation studies that found a direct involvement of these cortical areas in patients affected by visual neglect ([24], [54]–[58]). Our findings are consistent with the hypothesis that a behavioral task coupled with brain stimulation over the crucial cortical circuits facilitates learning ([23], [25], [38], [59]).

We run a second experiment to control for stimulation-related effects that were independent from training and interference between task conditions. In particular, we sought to understand whether the decrement we found in the OD task between baseline and post-training in Experiment 1 was related to the combination of the behavioral training with stimulation. Results showed that participants who received multi-session offline tRNS over parietal lobes performed significantly better on the OD task, while hMT+ and sham control groups did not improve. These results indicate that parietal stimulation can exert a beneficial effect even when delivered offline, and across multiple sessions at rest, on cortical areas that are crucial for a specific task ([60]). This effect is particularly interesting for its clear applicability to the pathological population, given the ease of use. In fact, stimulation alone would likely increase the compliance to treatment from stroke and other neurological patients ([61]–[63]).

In conclusion, we found clear opposite effects on the OD task on Day 5 relative to Day 1 between Experiment 1 and 2. We found a *detrimental* effect on OD when subjects received parietal tRNS concurrent with a competing training task (TOJ), and a *beneficial* effect on OD when subjects received stimulation while at rest. These results indicate that it is the ‘interference condition’ (Experiment 1) in which subjects are presented with task-relevant and task-irrelevant visual inputs that activates dual attentional mechanisms that promote the selection of relevant information while suppressing irrelevant ones. This mechanism is strongly activated when subjects are asked to detect the task-relevant vs irrelevant input within the same visual stimulus, as in our paradigm, and our data indicate a clear causal role of the parietal lobes in inhibitory mechanisms during learning. Overall, our results show that the parietal lobe is a central hub in gating and selecting crucial stimuli while inhibiting irrelevant information ([25], [49], [50]). Importantly, under conditions without (visuo-attentional) conflict, stimulation over the parietal lobes at rest can facilitate learning. Further investigation is needed to better understand how this protocol could be adapted for the healthy and the neurological population. Finally, our results also indicate that poor task selection in combination with tES could hamper the stimulation effect. While oftentimes the assumption is that a targeted transcranial electrical stimulation intervention might *enhance* a specific behavior, one should also consider that other variables might conflict and exert detrimental effect opposite to the expected outcome ([64]).

## Conclusions

Our neuromodulatory study demonstrates that the parietal lobe plays a crucial role in inhibitory processes during visual learning and that selective attention helps prioritize information while actively suppressing irrelevant features. Importantly, we show that tRNS can promote learning at rest without training, a procedure that could easily be implemented in the less compliant population such as neurological or psychiatric patients.

## Acknowledgements

L.B is supported by the National Institutes of Health (R01 AG060981-01), the Alzheimer’s Drug Discovery Foundation (ADDF), and The Blavatnik Family Foundation (Blavatnik Sensory Disorders Research Award).

## References

[1] L. Pessoa, S. Kastner, and L. G. Ungerleider, “Neuroimaging Studies of Attention: From Modulation of Sensory Processing to Top-Down Control,” 2003. Accessed: Jan. 25, 2022. [Online]. Available: https://www.jneurosci.org/content/jneuro/23/10/3990.full.pdf

[2] Y. Sasaki, J. E. Nanez, and T. Watanabe, “Advances in visual perceptual learning and plasticity.,” Nat Rev Neurosci, vol. 11, no. 1, pp. 53–60, Jan. 2010, doi: 10.1038/nrn2737.

[3] W. Li, V. Piëch, and C. D. Gilbert, “Perceptual learning and top-down influences in primary visual cortex.,” Nat Neurosci, vol. 7, no. 6, pp. 651–657, 2004.

[4] R. E. Crist, W. Li, and C. D. Gilbert, “Learning to see: experience and attention in primary visual cortex,” Nature Neuroscience, vol. 4, no. 5, pp. 519–525, May 2001, doi: 10.1038/87470.

[5] I. Mukai, D. Kim, M. Fukunaga, S. Japee, S. Marrett, and L. G. Ungerleider, “Behavioral/Systems/Cognitive Activations in Visual and Attention-Related Areas Predict and Correlate with the Degree of Perceptual Learning,” 2007, doi: 10.1523/JNEUROSCI.3002-07.2007.

[6] M. Ahissar and S. Hochstein, “The reverse hierarchy theory of visual perceptual learning,” Trends in Cognitive Sciences, vol. 8, no. 10, pp. 457–464, Oct. 2004, doi: 10.1016/j.tics.2004.08.011.

[7] P. R. Roelfsema, A. van Ooyen, and T. Watanabe, “Perceptual learning rules based on reinforcers and attention,” Trends in Cognitive Sciences, vol. 14, no. 2, pp. 64–71, Feb. 2010, doi: 10.1016/J.TICS.2009.11.005.

[8] M. Ahissar and S. Hochstein, “Attentional control of early perceptual learning.,” Proceedings of the National Academy of Sciences, vol. 90, no. 12, pp. 5718–5722, 1993.

[9] S. Spadone et al., “Dynamic reorganization of human resting-state networks during visuospatial attention,” vol. 112, no. 26, 2015, doi: 10.1073/pnas.1415439112.

[10] C. M. Lewis, A. Baldassarre, G. Committeri, and G. Luca, “Learning sculpts the spontaneous activity of the resting human brain,” pp. 1–6, 2009.

[11] S. Hochstein and M. Ahissar, “View from the Top: Hierarchies and Reverse Hierarchies Review,” vol. 36, no. 3, pp. 791–804, 2002.

[12] Z. L. Lu and B. A. Dosher, “Perceptual learning retunes the perceptual template in foveal orientation identification,” Journal of Vision, vol. 4, no. 1, pp. 5–5, Jan. 2004, doi: 10.1167/4.1.5.

[13] R. W. Li, D. M. Levi, and S. A. Klein, “Perceptual learning improves efficiency by re-tuning the decision ‘template’ for position discrimination,” NATURE NEUROSCIENCE, vol. 7, no. 2, 2004, doi: 10.1038/nn1183.

[14] I. Fine and R. A. Jacobs, “Comparing perceptual learning across tasks: A review,” Journal of Vision, vol. 2, no. 2, pp. 5–5, Apr. 2002, doi: 10.1167/2.2.5.

[15] A. Gazzaley, J. W. Cooney, K. Mcevoy, R. T. Knight, and M. D. ’ Esposito, “Top-down Enhancement and Suppression of the Magnitude and Speed of Neural Activity”, Accessed: Aug. 04, 2022. [Online]. Available: http://direct.mit.edu/jocn/article-pdf/17/3/507/1935119/0898929053279522.pdf

[16] T. Womelsdorf et al., “Modulation of neuronal interactions through neuronal synchronization,” Science (1979), vol. 316, no. 5831, pp. 1609–1612, Jun. 2007, doi: 10.1126/SCIENCE.1139597/SUPPL_FILE/WOMELSDORF-SOM.PDF.

[17] J. Duncan, S. Martens, and R. Ward, “Restricted attentional capacity within but not between sensory modalities,” Nature, vol. 387, no. 6635, pp. 808–810, Jun. 1997, doi: 10.1038/42947.

[18] Y. Sasaki and T. Watanabe, “Perceptual learning without perception,” vol. 413, no. October, 2001.

[19] Y. Tsushima, A. R. Seitz, and T. Watanabe, “Task-irrelevant learning occurs only when the irrelevant feature is weak,” Current Biology, vol. 18, no. 12, pp. R516–R517, Jun. 2008, doi: 10.1016/J.CUB.2008.04.029.

[20] R. Desimone and J. Duncan, “NEURAL MECHANISMS OF SELECTIVE VISUAL ATTENTION,” Annu. Rev. Neurosci, vol. 18, pp. 193–222, 1995, Accessed: Aug. 04, 2022. [Online]. Available: www.annualreviews.org/aronline

[21] J. H. Reynolds and D. J. Heeger, “The Normalization Model of Attention,” Neuron, vol. 61, no. 2, pp. 168–185, Jan. 2009, doi: 10.1016/J.NEURON.2009.01.002.

[22] F. Herpich, M. D. Melnick, S. Agosta, K. R. Huxlin, D. Tadin, and L. Battelli, “Boosting Learning Efficacy with Noninvasive Brain Stimulation in Intact and Brain-Damaged Humans,” The Journal of Neuroscience, vol. 39, no. 28, pp. 5551 LP – 5561, Jul. 2019, doi: 10.1523/JNEUROSCI.3248-18.2019.

[23] S. C. Tyler, F. Contò, and L. Battelli, “Rapid Improvement on a Temporal Attention Task within a Single Session of High-frequency Transcranial Random Noise Stimulation,” Journal of Cognitive Neuroscience, vol. 30, no. 5, pp. 656–666, May 2018, doi: 10.1162/jocn_a_01235.

[24] F. Contò, G. Edwards, S. Tyler, D. Parrott, E. Grossman, and L. Battelli, “Attention network modulation via trns correlates with attention gain,” Elife, vol. 10, Nov. 2021, doi: 10.7554/ELIFE.63782.

[25] G. Campana, R. Camilleri, A. Pavan, A. Veronese, and G. lo Giudice, “Improving visual functions in adult amblyopia with combined perceptual training and transcranial random noise stimulation (tRNS): a pilot study.,” Front Psychol, vol. 5, p. 1402, 2014, doi: 10.3389/fpsyg.2014.01402.

[26] G. Campana, R. Camilleri, B. Moret, F. Ghin, and A. Pavan, “Opposite effects of high- and low-frequency transcranial random noise stimulation probed with visual motion adaptation,” Scientific Reports, vol. 6, no. 1, p. 38919, Dec. 2016, doi: 10.1038/srep38919.

[27] M. A. Nitsche and W. Paulus, “Excitability changes induced in the human motor cortex by weak transcranial direct current stimulation,” pp. 633–639, 2000.

[28] M. A. Nitsche et al., “Transcranial direct current stimulation: State of the art 2008,” Brain Stimulation, vol. 1, no. 3, pp. 206–223, Jul. 2008, doi: 10.1016/j.brs.2008.06.004.

[29] M. A. Nitsche, M. S. Nitsche, C. C. Klein, F. Tergau, J. C. Rothwell, and W. Paulus, “Level of action of cathodal DC polarisation induced inhibition of the human motor cortex,” Clinical Neurophysiology, vol. 114, no. 4, pp. 600–604, Apr. 2003, doi: 10.1016/S1388-2457(02)00412-1.

[30] L. Chaieb, W. Paulus, and A. Antal, “Evaluating Aftereffects of Short-Duration Transcranial Random Noise Stimulation on Cortical Excitability,” vol. 2011, 2011, doi: 10.1155/2011/105927.

[31] L. Chaieb, W. Paulus, and A. Antal, “Evaluating aftereffects of short-duration transcranial random noise stimulation on cortical excitability,” Neural Plasticity, vol. 2011, 2011.

[32] F. Moss, L. M. Ward, and W. G. Sannita, “Stochastic resonance and sensory information processing: A tutorial and review of application,” Clinical Neurophysiology, vol. 115, no. 2. pp. 267–281, 2004. doi: 10.1016/j.clinph.2003.09.014.

[33] O. van der Groen and N. Wenderoth, “Transcranial Random Noise Stimulation of Visual Cortex: Stochastic Resonance Enhances Central Mechanisms of Perception.,” The Journal of neuroscience, vol. 36, no. 19, pp. 5289–5298, 2016.

[34] O. van der Groen, M. F. Tang, N. Wenderoth, and J. B. Mattingley, “Stochastic resonance enhances the rate of evidence accumulation during combined brain stimulation and perceptual decision-making,” PLOS Computational Biology, vol. 14, no. 7, p. e1006301, Jul. 2018, doi: 10.1371/journal.pcbi.1006301.

[35] A. Pavan, F. Ghin, A. Contillo, C. Milesi, G. Campana, and G. Mather, “Modulatory mechanisms underlying high-frequency transcranial random noise stimulation (hf-tRNS): A combined stochastic resonance and equivalent noise approach,” Brain Stimulation, vol. 12, no. 4, pp. 967–977, Jul. 2019, doi: 10.1016/J.BRS.2019.02.018.

[36] F. Herpich, F. Contò, M. van Koningsbruggen, and L. Battelli, “Modulating the excitability of the visual cortex using a stimulation priming paradigm,” Neuropsychologia, vol. 119, pp. 165–171, Oct. 2018, doi: 10.1016/J.NEUROPSYCHOLOGIA.2018.08.009.

[37] M. Cappelletti et al., “Transfer of Cognitive Training across Magnitude Dimensions Achieved with Concurrent Brain Stimulation of the Parietal Lobe,” vol. 33, no. 37, pp. 14899–14907, 2013, doi: 10.1523/JNEUROSCI.1692-13.2013.

[38] L. Battelli, A. Pascual-Leone, and P. Cavanagh, “The ‘when’ pathway of the right parietal lobe,” Trends in Cognitive Sciences, vol. 11, no. 5, pp. 204–210, May 2007, doi: 10.1016/J.TICS.2007.03.001.

[39] F. A. J. Verstraten, P. Cavanagh, and A. T. Labianca, “Limits of attentive tracking reveal temporal properties of attention,” Vision Research, vol. 40, no. 26, pp. 3651–3664, Dec. 2000, doi: 10.1016/S0042-6989(00)00213-3.

[40] D. H. Brainard and S. Vision, “The psychophysics toolbox,” Spat Vis, vol. 10, no. 4, pp. 433–436, 1997.

[41] D. G. Pelli and S. Vision, “The VideoToolbox software for visual psychophysics: Transforming numbers into movies,” Spat Vis, vol. 10, pp. 437–442, 1997.

[42] A. Thielscher, A. Antunes, and G. B. Saturnino, “Field modeling for transcranial magnetic stimulation: A useful tool to understand the physiological effects of TMS?,” in 2015 37th Annual International Conference of the IEEE Engineering in Medicine and Biology Society (EMBC), Aug. 2015, pp. 222–225. doi: 10.1109/EMBC.2015.7318340.

[43] G. B. Saturnino, H. R. Siebner, A. Thielscher, and K. H. Madsen, “Accessibility of cortical regions to focal TES: Dependence on spatial position, safety, and practical constraints,” Neuroimage, vol. 203, Dec. 2019, doi: 10.1016/J.NEUROIMAGE.2019.116183.

[44] B. A. Dosher and Z. L. Lu, “Perceptual learning reflects external noise filtering and internal noise reduction through channel reweighting,” Proc Natl Acad Sci U S A, vol. 95, no. 23, pp. 13988–13993, Nov. 1998, doi: 10.1073/PNAS.95.23.13988.

[45] J. H. Reynolds, L. Chelazzi, and R. Desimone, “Competitive Mechanisms Subserve Attention in Macaque Areas V2 and V4,” Journal of Neuroscience, vol. 19, no. 5, pp. 1736–1753, Mar. 1999, doi: 10.1523/JNEUROSCI.19-05-01736.1999.

[46] S. Kastner and L. G. Ungerleider, “The neural basis of biased competition in human visual cortex,” Neuropsychologia, vol. 39, pp. 1263–1276, 2001, Accessed: May 24, 2022. [Online]. Available: www.elsevier.com/locate/neuropsychologia

[47] S. Kastner, M. A. Pinsk, P. de Weerd, R. Desimone, and L. G. Ungerleider, “Increased activity in human visual cortex during directed attention in the absence of visual stimulation,” Neuron, vol. 22, no. 4, pp. 751–761, 1999, doi: 10.1016/S0896-6273(00)80734-5.

[48] C. L. E. Paffen, F. A. J. Verstraten, and Z. Vidnyánszky, “Attention-based perceptual learning increases binocular rivalry suppression of irrelevant visual features,” Journal of Vision, vol. 8, no. 4, pp. 25–25, Apr. 2008, doi: 10.1167/8.4.25.

[49] Z. Vidnyánszky and W. Sohn, “Learning to suppress task-irrelevant visual stimuli with attention,” Vision Research, vol. 45, no. 6, pp. 677–685, Mar. 2005, doi: 10.1016/J.VISRES.2004.10.009.

[50] U. Herwig, P. Satrapi, and C. Schönfeldt-Lecuona, “Using the International 10-20 EEG System for Positioning of Transcranial Magnetic Stimulation,” Brain Topography 2003 16:2, vol. 16, no. 2, pp. 95–99, Dec. 2003, doi: 10.1023/B:BRAT.0000006333.93597.9D.

[51] M. Okamoto et al., “Three-dimensional probabilistic anatomical cranio-cerebral correlation via the international 10–20 system oriented for transcranial functional brain mapping,” Neuroimage, vol. 21, no. 1, pp. 99–111, Jan. 2004, doi: 10.1016/J.NEUROIMAGE.2003.08.026.

[52] M. Corbetta and G. L. Shulman, “Spatial Neglect and Attention Networks,” *http://dx.doi.org/10.1146/annurev-neuro-061010-113731*, vol. 34, pp. 569–599, Jun. 2011, doi: 10.1146/ANNUREV-NEURO-061010-113731.

[53] L. Battelli et al., “Unilateral right parietal damage leads to bilateral deficit for high-level motion,” Neuron, vol. 32, no. 6, pp. 985–995, Dec. 2001, doi: 10.1016/S0896-6273(01)00536-0.

[54] L. Battelli, P. Cavanagh, P. Martini, and J. J. S. Barton, “Bilateral deficits of transient visual attention in right parietal patients,” Brain, vol. 126, no. 10, pp. 2164–2174, Oct. 2003, doi: 10.1093/BRAIN/AWG221.

[55] K. L. Roberts, J. K. L. Lau, M. Chechlacz, and G. W. Humphreys, “Cognitive Neuropsychology Spatial and temporal attention deficits following brain injury: A neuroanatomical decomposition of the temporal order judgement task,” 2012, doi: 10.1080/02643294.2012.722548.

[56] N. Dambeck et al., “Interhemispheric imbalance during visuospatial attention investigated by unilateral and bilateral TMS over human parietal cortices,” Brain Research, vol. 1072, no. 1, pp. 194–199, Feb. 2006, doi: 10.1016/J.BRAINRES.2005.05.075.

[57] S. Agosta et al., “The Pivotal Role of the Right Parietal Lobe in Temporal Attention,” Journal of Cognitive Neuroscience, vol. 29, no. 5, pp. 805–815, May 2017, doi: 10.1162/jocn_a_01086.

[58] A. Kiyonaga, J. P. Powers, Y. C. Chiu, and T. Egner, “Hemisphere-specific Parietal Contributions to the Interplay between Working Memory and Attention,” Journal of Cognitive Neuroscience, vol. 33, no. 8, pp. 1428–1441, Jul. 2021, doi: 10.1162/JOCN_A_01740.

[59] C. Pirulli, A. Fertonani, and C. Miniussi, “The role of timing in the induction of neuromodulation in perceptual learning by transcranial electric stimulation,” Brain Stimul, vol. 6, no. 4, pp. 683–689, 2013.

[60] C. I. Fernandez-Lazaro et al., “Adherence to treatment and related factors among patients with chronic conditions in primary care: A cross-sectional study,” BMC Family Practice, vol. 20, no. 1, pp. 1–12, Sep. 2019, doi: 10.1186/S12875-019-1019-3/TABLES/4.

[61] M. J. ElsnerB, “Cochrane Library Cochrane Database of Systematic Reviews Transcranial direct current stimulation (tDCS) for improving activities of daily living, and physical and cognitive functioning, in people aer stroke (Review) Transcranial direct current stimulation (tDCS) for improving activities of daily living, and physical and cognitive functioning, in people aer stroke (Review),” 2016, doi: 10.1002/14651858.CD009645.pub3.

[62] C. D. Solomons and V. Shanmugasundaram, “A review of transcranial electrical stimulation methods in stroke rehabilitation,” Neurol India, vol. 67, no. 2, p. 417, Mar. 2019, doi: 10.4103/0028-3886.258057.

[63] M. R. Krause, I. D. 1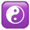, P. G. Vieira, J.-P. Thivierge, and C. C. Pack, “Brain stimulation competes with ongoing oscillations for control of spike timing in the primate brain,” 2022, doi: 10.1371/journal.pbio.3001650.

